# Growth Phase Dependent Chromosome Condensation and H-NS Protein Redistribution in *E. coli* Under Osmotic Stress

**DOI:** 10.1101/559138

**Authors:** Nafiseh Rafiei, Martha Cordova, William Wiley Navarre, Joshua N. Milstein

## Abstract

The heat-stable nucleoid-structuring (H-NS) protein is a global transcriptional regulator implicated in coordinating the expression of over 200 genes in *E. coli* bacteria. We have applied super-resolved microscopy to quantify the intracellular, spatial reorganization of H-NS in response to osmotic stress. We find that H-NS shows a growth phase dependent response to hyperosmotic shock. In stationary phase, H-NS detaches from a tightly compacted bacterial chromosome and is excluded from the nucleoid volume over an extended period of time. This behaviour is absent during rapid growth but may be induced by exposing the osmotically stressed culture to the DNA gyrase inhibitor, coumermycin. This compaction/H-NS exclusion phenomenon occurs in the presence of either potassium or sodium ions and is independent of the stress responsive sigma factor RpoS, or the H-NS paralog StpA.

## INTRODUCTION

Bacteria are under constant pressure to adapt and flourish within unstable and often unpredictable environments. One of the more effective means bacteria have for adapting to environmental stressors is by acquiring genetic traits through the mechanism of horizontal gene transfer (HGT). Expression of most horizontally acquired genes, however, will simply incur a metabolic cost or worse to the host placing the cells at a competitive disadvantage. Bacteria have therefore evolved to mitigate the costs of HGT with many enteric bacteria, such as *E. coli* and *Salmonella*, utilizing the heat-stable nucleoid-structuring (H-NS) protein. H-NS acts as a global transcriptional silencer of foreign (or xenogeneic) DNA acquired through HGT, primarily by binding to adenine and thymine rich regions of the chromosome^1^. H-NS has been implicated in coordinating a range of bacterial stress responses such as an adaptation to changes in pH, temperature and osmolarity^2^. Because of these observations, for decades H-NS was thought to primarily regulate the stress response of enterobacteria, until its role in silencing xenogeneic DNA was identified^3^. It is now generally appreciated that H-NS, like a eukaryotic histone, is a general factor that has evolved many regulatory roles including controlling the expression of AT-rich genes that are transcriptionally responsive to a variety of conditions including stress.

H-NS polymerizes on DNA to both block RNAP access to promoters and likely also to trigger RNAP stalls during its translocation/elongation along the DNA template^4-6^. As a consequence of its function, it also acts as a ‘domainin’ that prevents the diffusion of supercoils generated during replication and transcription along the DNA molecule, instead keeping such supercoils trapped to a local region of the chromosome^7-9^. Indeed the ability of H-NS to trap supercoils was identified in early studies on the biochemical properties of the molecule^10^. One reason H-NS may contribute to RNAP stalling is by preventing the free diffusion of positive supercoils in front of the translocating polymerase.

H-NS has been implicated in the regulation of the *E. coli* and *Salmonella* osmotic response both globally and specifically by several independent studies. H-NS is a direct negative regulator of the *proV* locus (where it was also named OsmZ in early studies)^11^. Globally, H-NS affects the expression of RpoS (σ^38^ or σ^S^) the RNAP sigma subunit that mediates enterobacterial responses to a variety of stressors including starvation and osmolarity^12,13^.

Supercoiling tension along any stretch of DNA is a combination of topoisomerase activity (stable changes to linking number), DNA binding proteins (domainins), and helicases/polymerases, which transiently cause supercoiling stress via the translocational unwinding of the DNA template. The findings that H-NS constrains supercoils, that H-NS negatively regulates the expression of several osmotically induced genes, and that high osmolarity alters the overall average negative supercoiling of several genes and plasmid reporters has led many to speculate that H-NS controls supercoiling in the cell. However, it is unclear whether H-NS binding is affected by DNA supercoiling, whether supercoiling is affected by H-NS directly or indirectly, or whether external triggers such as osmolarity or temperature can modulate the ability of H-NS to affect DNA supercoiling. These problems arise from the paucity of tools with sufficient resolution to dissect how H-NS responds *in vivo* to perturbations in environmental conditions.

In this study, we have focused on the spatio-temporal distribution of H-NS binding along the *E. coli* chromosome, in response to osmotic stress, and at different phases of growth (both exponential and early stationary phase). Previously, it was found that several highly expressed nucleoid associated proteins (NAPs) like H-NS distribute in a more or less random fashion about the nucleoid volume^14^. We should comment, since it is often incorrectly cited, that while H-NS was reported to cluster into foci within *E. coli*^14^, this observation proved to be an artefact of the fluorescent label mEos2^15^. Mutations to the dimerization domain of the label, resulting in the highly monomeric mEos3.2^16^, suppress the observed clustering resulting in the random patterning of H-NS throughout the nucleoid.

We have observed a pronounced, almost immediate, spatial redistribution of H-NS following osmotic shock during the stationary phase of growth. A rapid increase in osmolarity causes the bacterial chromosome to tightly condense, at which point H-NS apparently detaches from the chromosome and migrates toward the periphery of the cell. This behaviour is not observed in exponential phase under the same stress conditions, rather, H-NS remains distributed throughout the nucleoid despite a slight compaction of the chromosome. If, however, we subject exponentially replicating cells to the DNA gyrase inhibitor coumermycin^17-19^ during osmotic shock we observe both an increased compaction of the chromosome and an apparent detachment/exclusion of H-NS from the nucleoid volume, similar to what was observed in stationary phase. Dissociation of H-NS from the chromosome was also measured in chromatin immunoprecipitation assays. Notably the observed compaction and H-NS redistribution was independent of the central osmotic and stationary phase regulatory sigma factor, RpoS, or the nucleoid associated H-NS paralog StpA.

## MATERIAL AND METHODS

### Labelling H-NS with photoactivatable mEos3.2

A chimeric construct was created where H-NS was fused to the N-terminus of the monomeric, photoactivatable fluorescent protein mEos3.2. The coding region of H-NS from *E. coli* BW25113, fused to mEos3.2, was cloned upstream of the chloramphenicol resistance gene (*cat*) of plasmid pXG10^20^. This plasmid was then used as a PCR template to generate the fragment *hns*-mEos3.2-*cat* by PCR using Phusion High-Fidelity DNA Polymerase (Thermo Fisher) and primers SB039 (5’ - GCCGCTGGCGGGATTTTAAGCAAGTGCAATCTACAAAAGAGCTCTTTTTTGTGC GGTGCC) and SB40 (5’ - CCTCAACAAACCACCCCAATATAAGTTTGAGATTACTACACAACAGGAGTCCAA GCGAGC). The purified fragment was then used to replace the chromosomally encoded *hns* using the lambda red recombinase method described by Datsenko and Wanner^21^.The tagged chromosome was verified by Sanger Sequencing at the TCAG Sequencing Facility (Centre for Applied Genomics, Hospital for Sick Children) using primers EMC003 (5’ - GGTGTTATCCACGAAACGG) and EMC004 (5’ - CGTTAAATCTGGCACCAAAG).

### Cell fixation and sample preparation

We initially grew bacteria in M9 media (BioShop) supplemented with MgSO_4_ (2 mM), CaCl_2_ (0.1 mM), thiamine (0.01%), casamino acids (0.1%), and glucose (1%). Cells were shaken at 220 rpm at 37 ^°^C until they reached exponential (∼3 hours, OD_600_ ∼ 0.3) or stationary (∼ 6 hrs, OD_600_ ∼ 2) phase. We then exposed the cells to osmotic stress by adding KCl to the culture up to a final concentration of 300 mM (osmolarity ∼ 0.6 Osm/L). Substituting KCl for NaCl yielded no observable differences in cellular response. After exposure to the stress, we fixed the bacterial cells with 10x diluted formalin solution in H_2_O (Sigma-Aldrich, ∼37% formaldehyde stabilized with 10-15% methanol). In control experiments we found that fixation with methanol free formaldehyde (Thermo Fisher) did not have an effect on the observed results. The exposure time before fixation varied from 5 to 60 minutes for the time-course experiments, but for all other experiments the exposure time was set at 30 minutes. Cultures were incubated with formaldehyde while shaking at room temperature for 30 minutes. This was followed by 15 minutes of centrifugation at 1000 x g, discarding the supernatant and washing three times with 1 ml of PBS. For each washing step the pellet was first mixed with PBS for 5 minutes in an orbital shaker and then centrifuged for 10 minutes at 1000 x g at room temperature. After the final wash, the sample was ready to image or stored at 4 ^°^C to image within one week.

Samples were mounted by placing 3 μl of fixed cells atop a small 1.5% agarose pad. The agarose pad was then flipped over onto a coverslip that had been pre-cleaned with an initial 30 minutes of sonication in 3M KOH, rinsed in dH_2_O, then again sonicated for 30 minutes in ethanol (99%). Cells remained immobilized between the coverslip and the agarose pad. The coverslip was then affixed to the microscope slide with a 2.5 mm CoverWell spacer (Sigma-Aldrich) inserted in-between the two glass surfaces.

### Super-resolved microscopy

Microscopy was performed using an inverted Olympus IX-71 microscope equipped with a 60x oil-immersion TIRF objective (Olympus, APON 60XOTIRF, N.A. 1.49). Images were additionally magnified by a telescope inserted into the collection path of the microscope such that the effective pixel size was ∼73 nm, and collected by an electron multiplying charge-coupled device (EMCCD) camera (Andor, iXon3). All data acquisition was controlled by the open source microscopy program μManager^22^.

To obtain super-resolution radial fluctuation (SRRF) images, we conducted widefield fluorescence microscopy by imaging mEos3.2 in the green channel. We imaged the fluorescent protein by exciting with a low intensity (∼ 8 W/cm^2^) 488 nm laser (Coherent, OBIS) and transmitting the emission fluorescence using a 520/35 nm band-pass filter (Chroma). We acquired 100 frames for each ROI with a frame rate of 20 Hz. Subsequently, we applied SRRF to the image stack through an open source, GPU-enabled ImageJ plugin^23^. SRRF requires the user to define a ring radius for the underlying radiality analysis. A related technique “super-resolution quantitative image rating and reporting of error locations” (SQUIRREL), which is also available as an ImageJ plugin, can be used to evaluate the quality of the SRRF images^24^. Critical to achieving accurate images is the appropriate choice of the ring radius^23^. Therefore, we generated SRRF images from our raw image stack for various values of the ring radius, evaluated the resulting images with SQUIRREL, and chose the input value for the ring radius such that the error in the SRRF image was minimized.

### Nucleoid imaging

Fixed cells were stained with DAPI (3’, 6-diamidino-2-phynylindole) which enabled visualization of cellular DNA. We added this stain to fixed cultures to a final concentration of 0.1 µg/ml and incubated at room temperature for 5 minutes. Samples were then centrifuged at 5000 x g at room temperature for 1 minute, supernatant was discarded, and the pellets were washed by three subsequent resuspensions and centrifugations in 1 ml of fresh PBS. To image the DAPI-stained nucleoid of the bacterial cells, we employed an arc-lamp (X-cite series 120 Q), with an excitation band-pass filter 325/50 and an emission band-pass filter 447/60 inside the filter cube. For each image, a stack of 100 frames was acquired at a frame rate of 20 Hz.

### Image analysis and analytics

To quantify the spatial distribution of H-NS, cells were first manually segregated in ImageJ then analysed with custom MATLAB software. Cells that appeared to be dividing or that were roughly larger than 3 μm were removed from the statistical analysis. The principle axes of each cell were identified and the cells were then rotated through a linear transformation so their axes aligned. Cross sections were acquired by normalizing the axes of the cell from zero to one and plotting the intensity of a single pixel along the major principal axes.

Chromosome compaction was then quantified from fluorescence images by measuring the projected area occupied by the chromosome within the cell. Closed regions outlining the chromosome were automatically detected within ImageJ and the resulting areas calculated. Note, some cells displayed two distinct globular regions that we took to be two copies of the chromosome. For those cells, the area of each region was determined separately then added together. Finally, cell boundaries were acquired from bright-field images of the cells using ImageJ and used to normalize the area of the DNA containing nucleoid regions.

### Chromatin immunoprecipitation (ChIP)

Chromatin immunoprecipitation was carried out as described previously^25^. Briefly, *E. coli* BW25113 ∆*hns* was complemented with pWN426 (pHNS_HA_), which produces a functional variant of H-NS that carries a C-terminal HA (hemagglutinin) epitope tag that enables the protein to be immunoprecipitated with anti-HA antibodies^26^. This strain was grown in M9 media supplemented with MgSO4 (2 mM), CaCl2 (0.1 mM), thiamine (0.01%), casamino acids (0.1%), glucose (1%) and chloramphenicol (10 µg/ml). Overnight cultures were subcultured 1:50 in fresh M9 media and grown to exponential (OD_600_ ∼ 0.3) and stationary (OD_600_ ∼ 2) phase (matching the imaging conditions). Cells were then exposed to osmotic stress by adding KCl (300 mM) to the culture for 30 minutes. Coumermycin (final 5 µg/mL) was also added to a set of samples 30 min prior to the osmotic stress.

To crosslink protein to DNA from the different samples (50 mL each), formaldehyde was added to a final concentration of 1% for 15 min at room temperature before quenching with 1.25 mM glycine for 10 min. Cells were washed twice with cold PBS and sonicated to generate chromosomal fragments of average size ∼500 bp. Lysates were cleared by centrifugation and precipitated with anti-HA antibody (clone HA-7; Bioshop) using agarose protein G beads (Calbiochem) as previously described^25^. The samples were incubated at 65 °C for 5 h to break the DNA-protein crosslinks.

DNA fragments that co-precipitated with H-NS_HA_ were quantified by real-time quantitative PCR using SsoFast EvaGreen Supermix (Bio-Rad) according to the manufacturer’s instructions and using gene-specific primers: forward 5-ACACTGTTAACCGCCAGGAAGAC A-3 and reverse 5-GGATGAAAGCAAAGCGCAAGCAGA-3 for *bglG*; forward 5-AGTTCCGTGCAGGAAGAGAACCTT-3 and reverse 5-TGGTTACG TCGCTTTCGGCTTACT-3 for *yjcF*; forward 5-AATATTTGGCGAGCATCCA CAGCG-3 and reverse 5-TTTACCCGAGCCGGATAATCCCAT-3 for *proV*; forward 5-GCAATCGACGCGATTCTTCCATCAAG-3 and reverse 5-GCAGC GCGTTTAAATATGTCTCAGCC-3 for *xapR*.

## RESULTS

### In situ analysis of H-NS and nucleoid localization

We proceeded to visualize the spatial distribution of H-NS and the nucleoid using super-resolved radial fluctuations (SRRF) imaging^23^; an approach similar to techniques like super-resolution optical fluctuation imaging (SOFI)^27^ that make use of temporal correlations in an image stack to achieve super-resolved image reconstructions. SRRF applied to a conventional, wide-field image stack can obtain a resolution of approximately 100-150 nm, which provided sufficient detail for this study.

For the imaging, the chromosomal *hns* gene was engineered with a 3’ fusion to the gene encoding the monomeric photo-activatable fluorescent protein mEos3.2 to generate a chimeric protein, H-NS-mEos. While we did not make use of the photoswitchable properties of mEos3.2 here, it’s proven ability to remain truly monomeric when fused to H-NS makes it an ideal choice for this study^15^. In addition, our initial pilot studies utilized H-NS-mEos expressed from a low copy plasmid. However, we found that plasmid mediated expression of H-NS-mEos led to the observation of H-NS foci, that were likely caused by H-NS association with the expression plasmids themselves. Similar experiments run with a chromosomally encoded version of H-NS-mEos did not show these punctate foci. The nucleoid was imaged by staining with DAPI.

### Dynamics of the nucleoid in response to osmotic shock during growth and stationary phase

A previous study found that RNAP is displaced and the nucleoid condenses dramatically under osmotic shock when exponentially growing *E. coli* cells are treated with KCl^28^. We examined the dynamic response of the nucleoid to osmotic stress by fixing cells at a series of time points following induction of osmotic shock. **Figures 1A** and **1B**, respectively, show the absolute and fractional (relative to that of the cell) area of the nucleoid for cells in exponential growth or stationary phase. The analysis is of DAPI stained chromosomal DNA at 0, 5, 10, 20, 30, 45, and 60 minutes after the addition of 300 mM KCl.

**Figure 1.**
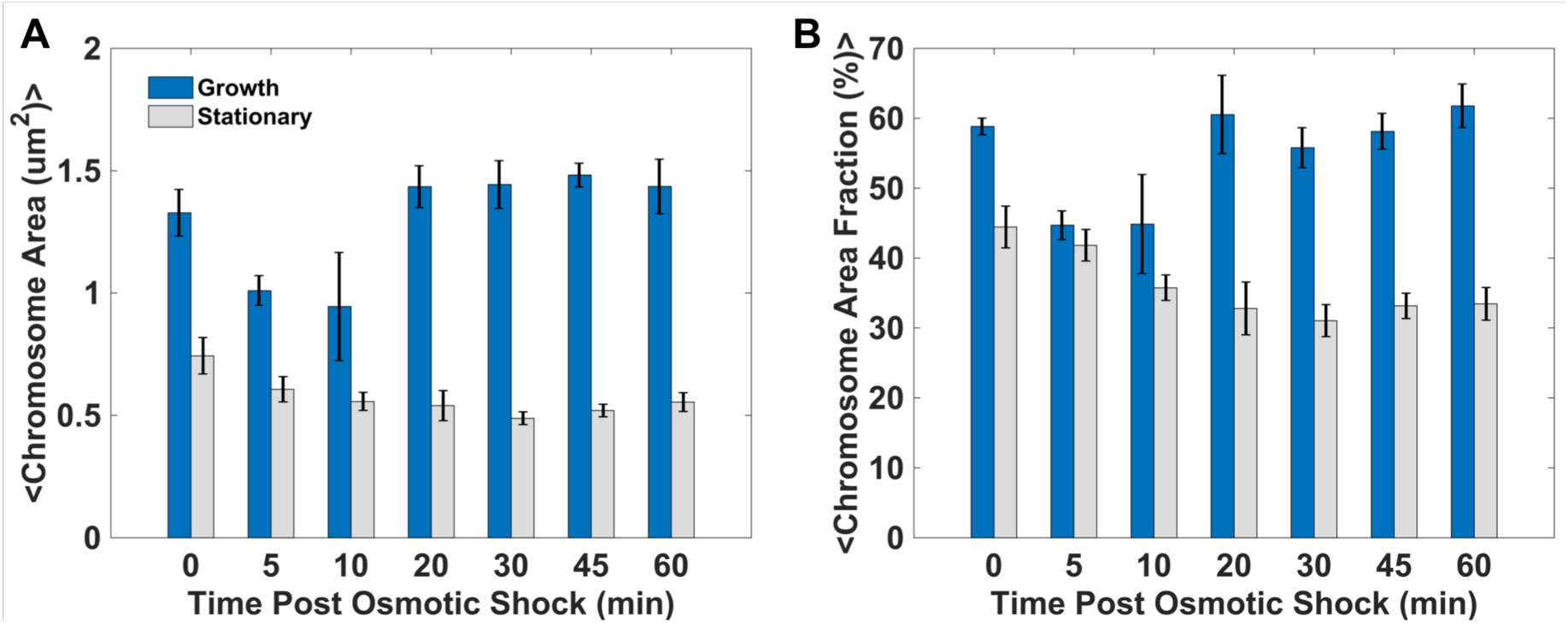
Dynamics of chromosome compaction following osmotic shock. **A)** Absolute, projected chromosome area and **B)** the area fraction (i.e., normalized to the projected area of the cell), for both exponential and stationary phase up to 60 minutes post osmotic shock. Error bars represent the standard deviation of the mean.

In rapidly growing cells the nucleoid shows significant condensation between 5 to 10 minutes, but by 20 minutes the images of the chromosome are difficult to differentiate from those taken before the changes in pre-induction. We presume this is due to rapid adaptation of the cells to osmotic stress. In stationary phase, the nucleoid is more compact pre-shock and condenses less dramatically than the nucleoid of cells in exponential growth after the addition of KCl. However, after the addition of KCl the nucleoid remains condensed and does not return to its pre-shock size after 60 minutes.

### H-NS is displaced from the stationary phase chromosome during osmotic shock

Concurrent with imaging the chromosome, we examined the localization of H-NS-mEos at various time points post osmotic shock in both rapidly growing (**Figure 2**) and stationary phase cells (**Figure 3**). In both figures, images are provided for 0, 5, 10, and 45 minutes post induction.

**Figure 2.**
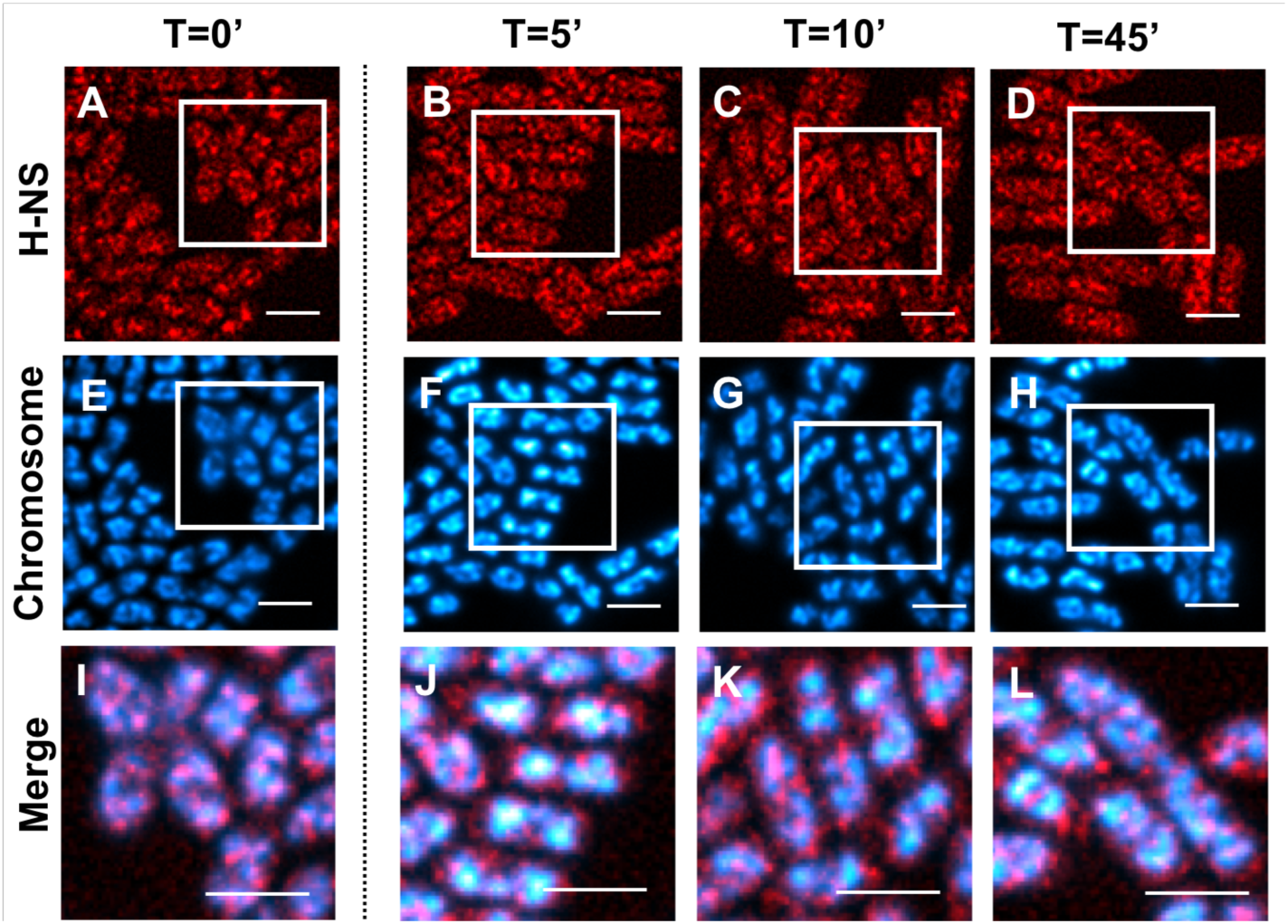
SRRF images of growth phase dynamic response to osmotic stress (300 mM KCl). From left to right: t=0, 5, 10, and 45 min post induction. **A-D)** H-NS, **E-H)** chromosome and **I-L)** merged image magnified to show detail (associated regions are indicated by square boxes in the images above). All scale bars are 2 μm.

**Figure 3.**
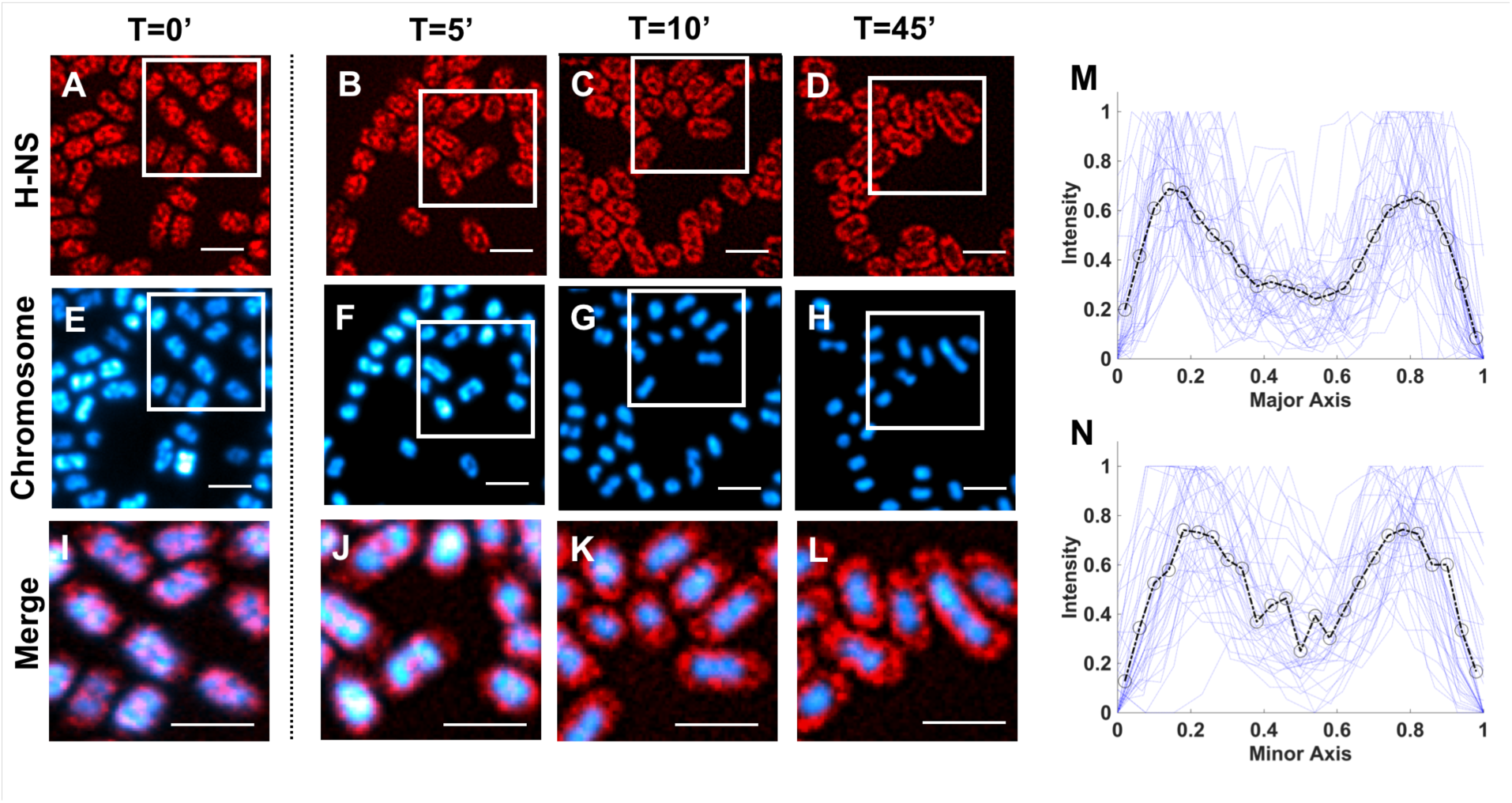
SRRF images of stationary phase dynamic response to osmotic stress (300 mM KCl). From left to right: t=0, 5, 10, and 45 min post induction. **A-D)** H-NS, **E-H)** chromosome and **I-L)** merged image magnified to show detail (associated regions are indicated by square boxes in the images above). All scale bars are 2 μm. **M)** Major and **N)** minor axis normalized intensity cross sections of H-NS distribution at 30 minutes post induction. Single bacteria intensity traces are shown with the average given by the dashed line (mean pixel values indicated by circles).

Post induction of osmotic shock, no noticeable response is observed in the spatial distribution of H-NS during the growth phase (**Figure 2**). However, during stationary phase (**Figure 3**), after 5 minutes H-NS can be seen to be reorganizing throughout the cell volume, forming a ring-like pattern that is fully established by 10 minutes post induction and maintained for at least 60 min, which is the longest period for which we measured. The chromosome, which is already more condensed in stationary phase than in exponential phase, continues to compact up until roughly 10 minutes post-induction. From the images, the osmotically shocked stationary-phase chromosome appears to be tightly condensed within the centre of each cell with the bulk of H-NS surrounding it.

### Coumermycin inhibition of DNA gyrase alters the nucleoid response to osmotic shock during rapid growth

Cells were then treated with 5 μg/ml coumermycin, an antibiotic which acts as a DNA gyrase inhibitor^17-19^, for 30 min and then imaged at 30 min post osmotic induction. No effect was seen in stationary phase; however, cells within exponential phase now display a spatial reorganization of H-NS similar to what was previously observed only in stationary phase (**Figure 4**). Once again, the bacterial chromosome is observed to be significantly more compact than pre-osmotic induction and H-NS is again seen to be excluded from the chromosomal volume, pushed toward the periphery of the bacterial cell. Note, this behaviour is not seen in exponentially growing cells treated with coumermycin alone.

**Figure 4.**
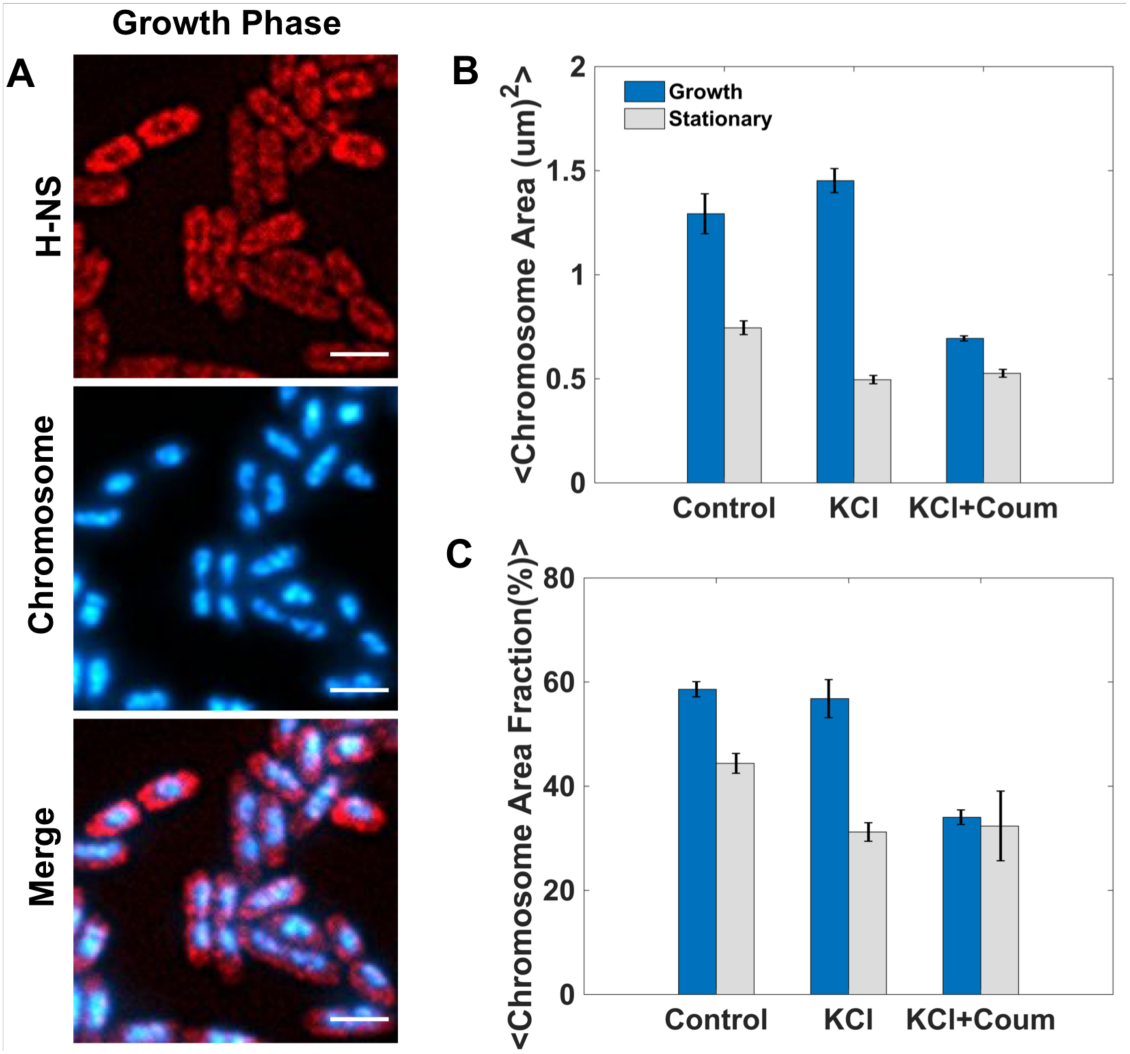
Analysis of coumermycin (5 µg/mL) treated cells. **A)** SRRF images in exponential growth phase 30 min post osmotic stress (300 mM KCl). From top to bottom: H-NS, chromosome, and the merged image. Scale bars are 2 μm. **B)** Comparison between exponential growth and stationary phase of the projected chromosome area and **C)** the area fraction for control, osmotic shock (KCl), and KCl + coumermycin exposed cells.

### Chromatin immunoprecipitation of H-NS during osmotic shock

A prior study examined the effect of transient K^+^ accumulation on the organization of the nucleoid and the association of RNAP^28^. Using cell fractionation, Cagliero *et al.* also noted that H-NS appeared to transiently and partially dissociate from the nucleoid before reassembling after osmoadaptation^28^. If H-NS does dissociate from the chromosome it should manifest as a decrease in DNA binding by chromatin immunoprecipitation assays. The alternative hypothesis is that while H-NS migrates to the periphery of the cell during chromosomal compaction, the H-NS bound DNA sequences remain associated with the protein and “loop” away from the bulk of the nucleoid, which might not be discernible by microscopy.

To test whether H-NS was indeed displaced from its bound loci during osmotic shock we performed chromatin immunoprecipitation on four known H-NS bound loci around the chromosome (*proV, xapR, yjcF* and *bglG*) under both stationary and rapid growth conditions before and after treatment with sodium chloride and/or coumermycin (**Figure 5**). Consistent with what was observed by microscopy, we find that H-NS association with its DNA targets (e.g., *proV*) is diminished in response to a rapid increase in osmolarity and this dissociation is particularly acute during stationary phase. Coumermycin on its own did not dramatically affect H-NS binding in either growth phase but dramatically potentiated the displacement of H-NS caused by an increase in osmolarity during exponential growth.

**Figure 5.**
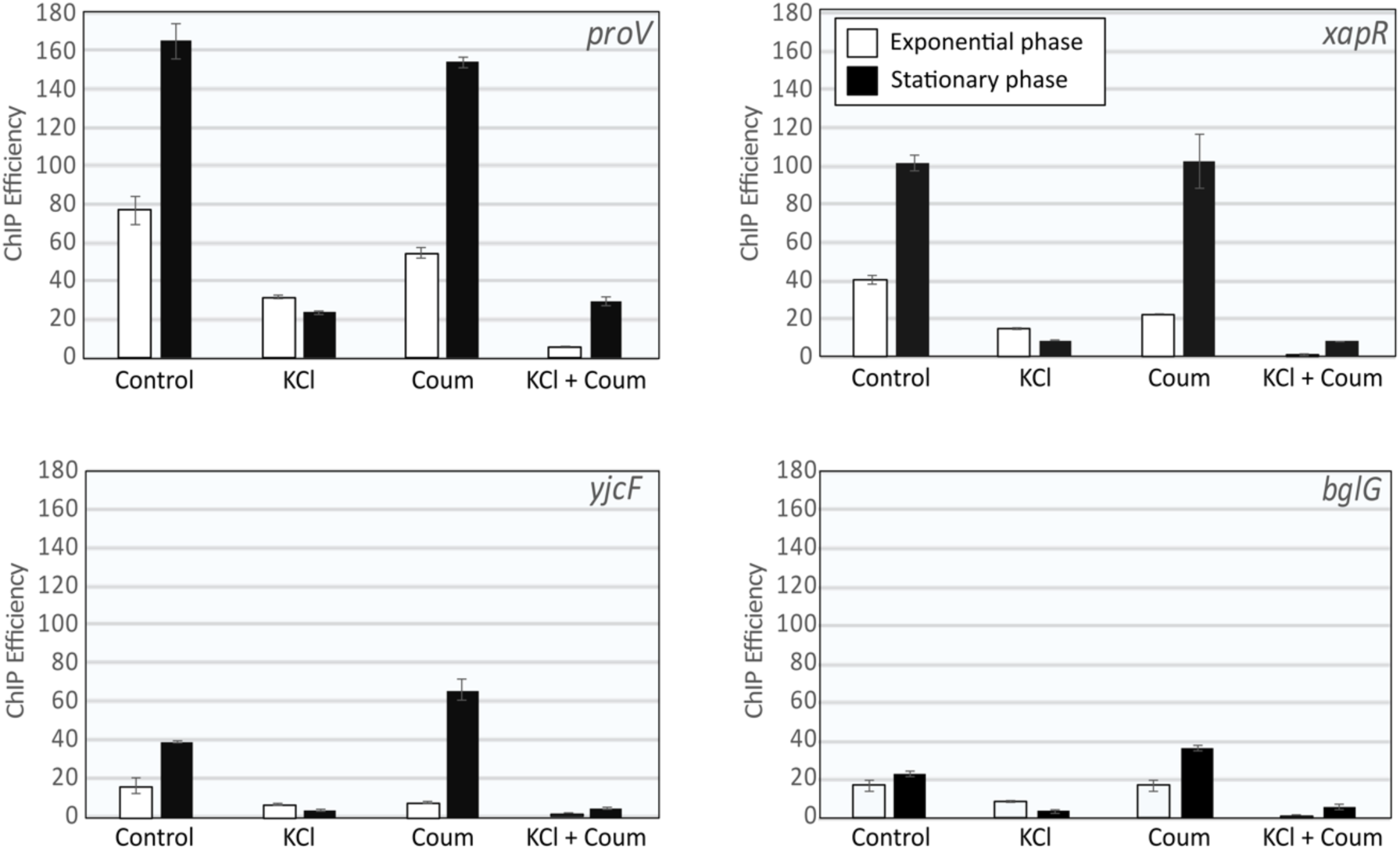
Chromatin immunoprecipitation of H-NS_HA_ at four H-NS bound loci (*proV, xapR, yjcF, bglG*) during osmotic shock (KCl), gyrase inhibition w/ coumermycin (Coum), or both. Cells, diluted in M9 media from overnight cultures, were grown to the indicated growth phase. 30 minutes after addition of KCl, coumermycin, or both, cells were fixed with formaldehyde and DNA-protein complexes were immunoprecipitated as described in the methods section. Enrichment of bound DNA was quantified by real-time PCR and normalized to the amount of DNA in the sample prior to immunoprecipitation.

### RpoS and StpA are not involved in the observed compaction of the nucleoid or expulsion of H-NS

We entertained the idea that H-NS might transiently be displaced from the bacterial chromosome during osmotic shock through competition from its paralog StpA. Notably, StpA demonstrates higher affinity for DNA *in vitro* in the presence of salt ^29,30^. To assess whether StpA plays any role in displacing H-NS from the nucleoid we visualized H-NS-mEos localization in an isogenic strain harbouring a deletion in *stpA*. We observed that H-NS was displaced from the chromosome similarly either in the absence or presence of StpA, indicating that competitive binding was not the root cause of H-NS depletion from the chromosome (data not shown).

RpoS is a central and pleiotropic regulator of the cellular response to osmotic stress^31^. We assessed whether a member of the RpoS regulon may be directly or indirectly involved in displacing H-NS from the chromosome under osmotic shock. However, we found that H-NS-mEos localization was similar between strains having or lacking functional RpoS (data not shown). This fact, along with the speed at which it occurs, suggests that H-NS displacement from the chromosome may be a direct consequence of the physical changes that occur inside the cell as a result of osmotic shock, and is not a downstream event that is controlled by regulatory factors within the cell.

## DISCUSSION

On average, the bacterial chromosome of *E. coli* is negatively supercoiled. However, the level of negative superhelicity varies throughout the growth phase with the overall level decreasing as the cell moves from exponential to stationary phase^32,33^. This transition is thought to be mediated primarily by changes in DNA gyrase activity and the altered binding of nucleoid associated proteins (NAPs). DNA gyrase induces negative supercoiling, which facilitates transcription by relieving torsional stress along double stranded DNA, while NAPs are conjectured to stabilize the level of local supercoiling throughout the chromosome^18,34-38^.

We have observed that during stationary phase, under conditions of osmotic stress, the *E. coli* chromosome tightly condenses and the nucleoid associated protein H-NS is actively excluded from the DNA containing volume of the cell, relocating toward the membrane periphery. While this behavior was observed in stationary phase, such gross spatial reorganization was not seen under identical conditions during rapid growth—despite the concurrent observation that the chromosome condenses following osmotic shock in both phases, although more so during stationary phase.

We were then able to reproduce a similar effect in the exponential phase upon addition of coumermycin, an antibiotic compound that inhibits DNA gyrase activity. Following exposure to coumermycin, the chromosomal DNA showed an increased level of compaction and H-NS was again excluded from the nucleoid volume, similar to what we had previously observed only in stationary phase. When we then repeated the coumermycin experiments in stationary phase, again under conditions of osmotic stress, we detected no observable changes. The chromosome remained significantly condensed with H-NS excluded from the nucleoid volume.

We provide a few possible explanations for our observations. In one model, osmotic stress induces the rapid condensation of the bacterial chromosome. This occurs in both exponential and stationary phase, though to different extents. In stationary phase, where the DNA already displays a reduced level of negative supercoiling pre-osmotic stress conditions, DNA condensation is so extreme that H-NS rapidly dissociates from the DNA. In exponential phase, this response is attenuated by the activity of DNA gyrase, which keeps the overall level of negative supercoiling elevated in preparation for higher levels of transcriptional activity than in stationary phase. The chromosome condenses, but to a limited extent, and H-NS is still able to remain bound. However, by interfering with DNA gyrase activity with coumermycin, the level of negative helicity is reduced and the chromosome is now able to tightly compact removing H-NS from the DNA.

Yet another model posits that high osmolarity in the cytoplasm reduces the affinity of H-NS for DNA and that large-scale H-NS dissociation releases previously trapped supercoils, which causes the compaction of DNA (once again, this would be more severe in stationary phase cells). This model suggests that the disruption of H-NS binding is the “proximal” event that occurs prior to chromosome compaction. The bacterial cell would then adapt via the production and accumulation of compatible solutes, which would ultimately restore homeostasis. Therefore, we remain unclear on the order; whether nucleoid compaction directly causes H-NS to dissociate or whether dissociation of H-NS leads to chromosomal compaction.

It is perhaps counterintuitive that relaxing negative supercoils, instead of causing an expansion, leads to a tighter compaction of the nucleoid, yet this phenomenon has been observed before by other groups^28,39^. Since supercoiling is known to affect the binding affinity of many proteins, it could be that a reduction in negative superhelicity causes other proteins to bind to and further condense the chromosomal DNA. Beyond simply being counterintuitive, a real complication to this model is that several groups have found osmotic stress to increase, not decrease, the level of negative supercoiling within the bacterial cells^11,40,41^. We note that these previous studies examined DNA topology using linkage assays or cruciform formation in plasmid reporters in actively growing cells or after extended periods of growth at high osmolarity, all of which differ from the specific conditions under which we observed chromosomal compaction. But if the extreme compaction of the chromosome, and subsequent expulsion of H-NS, is a result of increased negative supercoiling, then why do we observe this behaviour in stationary phase and not exponential unless we inhibit DNA gyrase? While we are unable to answer this question at present, the effect is quite apparent in our data and it suggests a simple, biophysical mechanism for responding to osmotic stress by preventing H-NS mediated gene silencing of particular stress response genes.

For instance, one pathway through which H-NS might coordinate an osmotic stress response is by regulating the stationary phase specific σ^S^ subunit of RNA polymerase. Encoded by the gene *rpoS*, σ^S^ is a global regulator of the stationary phase stress response of *E. coli.* Both σ^S^ and many σ^S^-regulated genes are induced under conditions of osmotic stress experienced in stationary phase^42,43^. H-NS, however, has been demonstrated to repress σ^S^ expression, which implies that derepression of the σ^S^-regulatory network by H-NS is necessary to trigger osmotic adaption. The dissociation of H-NS from the chromosomal DNA shortly after osmotic induction would enable the σ^S^ stress response network to activate. Note, we also performed these experiments in *rpoS* deficient cells, which had no effect on our observations; *rpoS* does not appear to play a role in condensing the chromosome or expelling H-NS from the nucleoid volume following osmotic stress in stationary phase.

Finally, we note that during our review of the literature there remain a number of apparently contradictory statements and views regarding supercoiling, osmolarity, and gene regulation. It is clear from our study and others that growth phase, source of osmotic stress, and length of exposure all play a role in what has been observed, which makes comparisons between studies challenging. Furthermore, it remains unclear how well plasmid linking numbers can serve as surrogate metrics for supercoiling-mediated phenomena that may be local. It also remains difficult to dissect what effects are bona fide regulatory responses to changes in supercoiling as opposed to adventitious secondary consequences caused by perturbations in nucleoid structure.

## ACKNOWLEDGEMENT

This work was funded by the Natural Sciences and Engineering Research Council of Canada [J.N.M., N.R.] (RG 418251), an Early Researcher Award from the Ministry of Research and Innovation [J.N.M., N.R.], and the Canadian Institutes of Health Research [W.W.N] (MOP-86683).

